# Molecular Time Capsules Enable Transcriptomic Recording in Living Cells

**DOI:** 10.1101/2023.10.12.562053

**Authors:** Mirae Parker, Jack Rubien, Dylan McCormick, Gene-Wei Li

## Abstract

Live-cell transcriptomic recording can help reveal hidden cellular states that precede phenotypic transformation. Here we demonstrate the use of protein-based encapsulation for preserving samples of cytoplasmic RNAs inside living cells. These molecular time capsules (MTCs) can be induced to create time-stamped transcriptome snapshots, preserve RNAs after cellular transitions, and enable retrospective investigations of gene expression programs that drive distinct developmental trajectories. MTCs also open the possibility to uncover transcriptomes in difficult-to-reach conditions.

---

Gene expression is dynamic and shapes future cellular states, but such temporal trajectories remain challenging to map at scale. Many important biological processes, such as cellular differentiation and disparate survivability under stress, are seeded by subpopulations of cells with distinct transcriptomic signatures^1–4^. Mapping these time-dependent trajectories requires capturing the transcriptome specific to the transient and founding population before a distinguishable phenotype has emerged. However, most methods to probe global gene expression necessitate immediate destruction of the cell, preventing longitudinal tracking of subpopulations. Although single-cell transcriptomics provide an opportunity to dissect populational heterogeneity^5,6^, they are also end-point measurements and must rely on inference methods, such as RNA velocity^7–10^, to postulate temporal dynamics.

Meanwhile, live-cell methods for monitoring gene expression remain limited in coverage, throughput, or temporal-resolution. Even with advanced multicolor fluorescence microscopy, there are insufficient channels to simultaneously monitor the entire genome, therefore necessitating prior knowledge on marker gene selection to investigate expression-to-phenotype trajectories^11–13^. Live-cell continual RNA extraction using force microscopy (e.g., Live-Seq^14^) can achieve full genome coverage and longitudinal tracking of single cells, though it currently has a limited throughput of 100s of cells and necessitates the adoption of specialized equipment. Recent breakthroughs in gene-expression recording by DNA-editing^15–20^ provide another promising avenue for interrogating time trajectories with a sequencing readout. These genome-based recording techniques currently require hours of active recording to accumulate dozens of events per lineage or are limited to recording a handful of pre-selected gene targets. We reasoned that a whole-transcriptome recording method that can be induced on-demand in a large population of cells could help elucidate the transitory expression changes that drive cellular differentiation. A tool with this capability could also enable the investigation of cell physiology in difficult-to-reach conditions, such as microbes in animal guts^21,22^.

To establish such a method, we reasoned that self-assembled protein capsules can be used to encapsulate and preserve a fraction of the cytoplasm in living cells (Fig. 1a). Capsule assembly can be controlled on-demand via an inducible promoter, creating a time-stamped record that is maintained in the cell. At a later time, such as when cells become phenotypically distinguishable or accessible, the capsules can be isolated via affinity purification and their content analyzed (Fig. 1b). We focus on their RNA content in this study, although these “molecular time capsules” (MTCs) could in principle be also used for proteomic or metabolomic studies.

**Fig 1.**
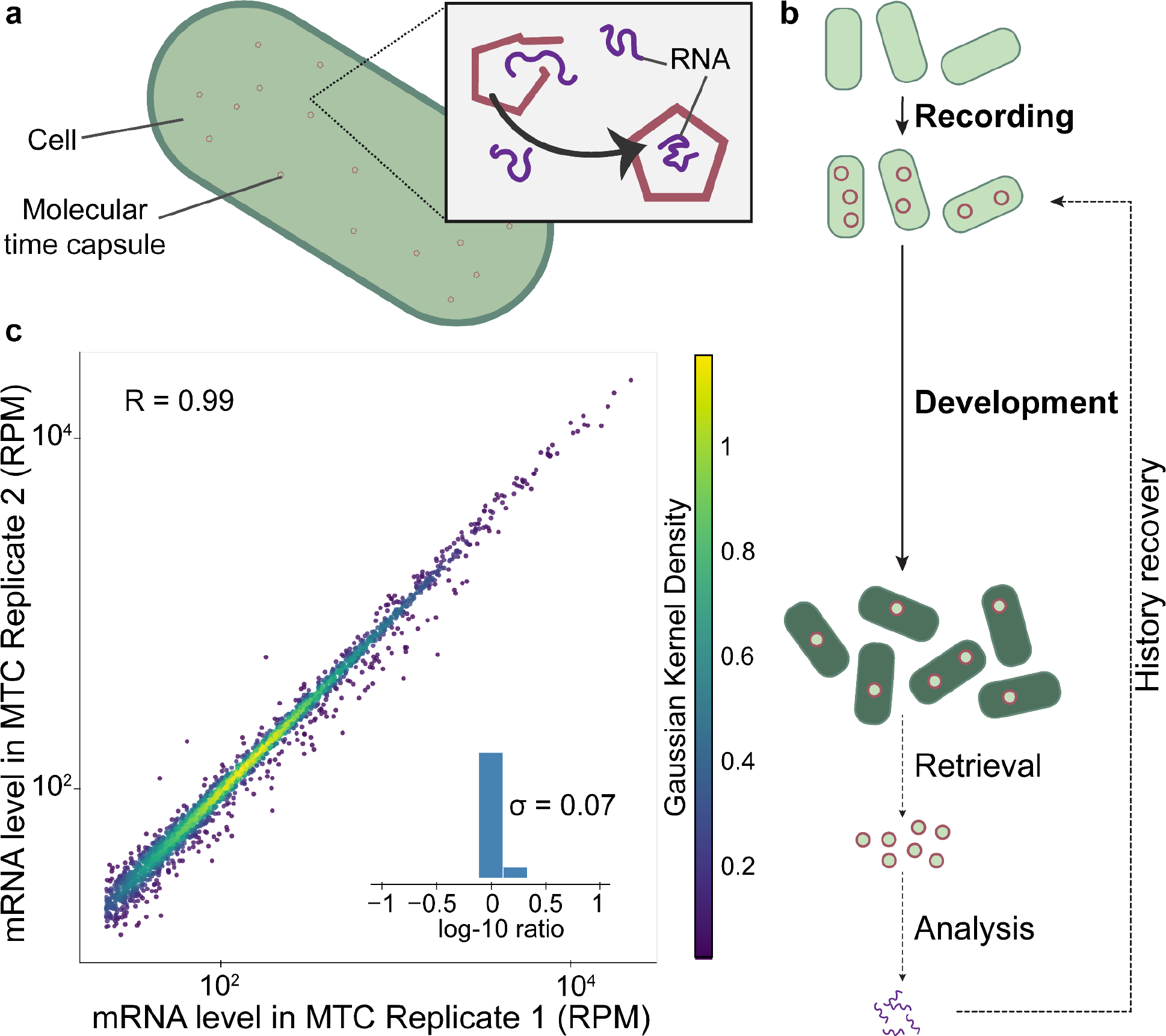
Molecular time capsules provide reproducible transcriptome recording in living cells. **a**, <u>M</u>olecular time <u>c</u>apsules (MTCs) are genetically encodable protein capsules that can self-assemble within living cells. During assembly, MTCs can encapsulate RNAs (see inset). **b**, MTCs capture and protect snapshots of transcriptomes for delayed retrieval and analysis. After a “recording” period, MTC production is shut off, and the cells are allowed to continue along their developmental trajectory. After the phenotype of interest has emerged, MTCs are retrieved by affinity purification and their encapsulated RNA is extracted. Sequencing analysis is performed on the RNAs, facilitating historical recovery of the transcriptomic record that was captured during the recording window. **c**, MTC-captured snapshots of the transcriptome across 2 biological replicates plotted in units of reads per million (RPM), where each dot corresponds to a gene with sufficient sequencing coverage. (Pearson correlation coefficient of log-transformed RPM values: R = 0.993, n = 2,302 genes, only genes with > 100 reads are considered). The inset plots the distribution of log-10 fold change between the two replicates. The standard deviation, σ, of the fold-change distribution = 0.07, or 1.2 fold.

Several criteria are required for faithful transcriptome recording by MTCs. First, the captured RNA content must be reproducible. Second, the retrieved RNA content must have minimal contamination from non-encapsulated RNAs, whether from the host cell or from other cells in the sample. Third, the RNA record must be stably preserved over time despite changes in the host transcriptome. Here we demonstrate that these criteria are met using a *de novo* designed protein capsule^23^(13.5-nm radius) expressed with a poly-histidine tag in *Escherichia coli*, with the recording timing controlled via an Isopropyl ß-D-1-thiogalactopyranoside (IPTG)-inducible promoter.

First, we assessed the reproducibility of MTC-based recording by comparing the encapsulated RNA content from three biological replicates. Poly-histidine-tagged MTCs were purified using a Ni-NTA column (Extended Figure 1) from exponential-phase cells that have been expressing capsule proteins steadily (for >10 generations). Expression of MTCs only slightly increases the population doubling time (from 20 minutes to 28.5 minutes) (Extended Figure 2). When expressed in this experiment, MTC transcripts represent 0.14% of all sequenced transcripts (transcripts per million (TPM) = 1350 +/- 230). Our purification and RNA extraction methods extract RNA specific to MTCs, as we are not able to recover a detectable amount of RNA from wild-type *E. coli* cells subjected to the same protocol.

Overall, the levels of encapsulated mRNAs, as measured by RNA-seq, agree across biological replicates on a gene-by-gene basis. The median Pearson correlation coefficient of log-transformed mRNA levels between replicates is R = 0.985 (Fig. 1c and Extended Figure 3). This reproducibility suggests that MTCs can facilitate differential expression analysis between cellular states. The mRNA levels in the total cell lysate also correlate with MTC samples, albeit less well compared to the reproducibility of MTC capture across biological replicates (median R = 0.784, Extended Figures 3 and 4). Interestingly, the majority of RNA fragments recovered from MTCs are short (< 200 nt), suggesting that MTC-based capture leverages the abundant RNA decay intermediates recently reported to dominate the transcriptome^24^. (Extended Figure 5).

We next demonstrated that MTC-encapsulated transcripts can be cleanly recovered without substantial contamination from non-encapsulated RNAs. To do so, we purified MTCs from a heterogeneous cell culture in which MTCs are only expressed in one subpopulation (*E. coli*, 50% of cells) but not in the other (*Bacillus subtilis*, 50% of cells) (Fig. 2a). These bacterial species have distinct genomes, facilitating the identification of which species a particular RNA originates from RNA-seq analysis.

**Fig 2.**
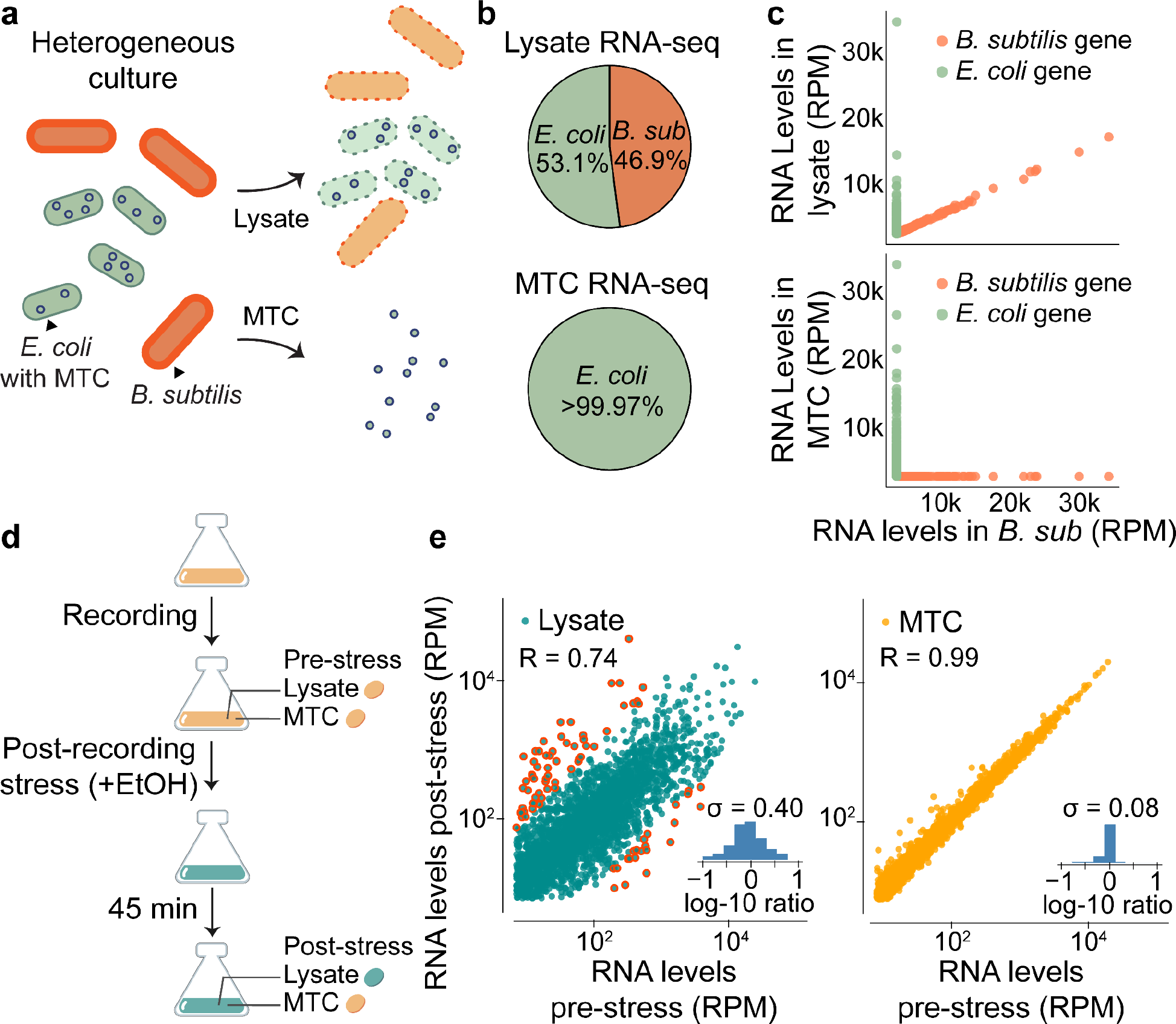
MTCs provide host-specific and stable storage. **a**, Total lysate and MTC samples are removed from a mixed “barnyard” sample consisting of both *B. subtilis* (orange) and MTC-containing *E. coli* (green). The MTC sample refers to MTCs purified from the mixed population. **b**, The percentage of reads in the MTC sample mapping to the *E. coli* genome is >99.97% (out of n = 5.2 x 10^6^ uniquely mapping reads), the mixed lysate sample has 53.1% (out of n = 3.2 x 10^6^ uniquely mapping reads) of reads uniquely mapping to *E. coli* genome. **c**, The gene-by-gene scatter plot in units of reads per million (RPM) for samples purified from pure *B. subtilis* lysate (x axis) versus the mixed lysate sample (top y-axis) and the MTC sample (bottom y-axis). *E. coli* genes are colored in green, whereas *B. subtilis* genes are colored in orange. **d**, MTC encapsulated RNA samples and lysate samples are collected from before and after a stressful treatment of 4% ethanol. **e**, The reads per million (RPM) of all genes with sufficient sequencing coverage are plotted, with each point corresponding to a separate gene. The scatter plot places a dot for each gene corresponding to its pre- (x) and post- (y)stress time-points of the lysate samples (left, blue) and the MTC samples (right, orange), along with accompanying insets showing the log-10 fold distribution between the pre- and post-stress samples. For the lysate samples, the Pearson correlation coefficient of log-transformed RPM values was R = 0.744, and the standard deviation (σ) of the log-10 fold-change distribution = 0.40, which corresponds to 2.5-fold (n = 2477 genes, only genes with more than 100 reads in both samples are considered.). Genes with greater than or equal to an order of magnitude change in expression between the pre- and post-stress samples are circled in red. For the pre- and post-stress MTC samples, the Pearson correlation coefficient of the log-transformed RPM values was R = 0.991, and the σ of the fold-change distribution = 0.08, which corresponds to 1.2-fold, (n = 2543 genes, only genes with more than 100 reads in both samples are considered.)

Although 46.9% of uniquely mapping reads from the heterogeneous lysate mapped to the *B. subtilis* genome, MTCs extracted from the same lysate contain almost exclusively *E. coli* RNAs (99.97%) (Fig. 2b). Every *B. subtilis* gene is depleted in MTCs (Fig. 2c). The level of *B. subtilis* mapping RNAs (0.03%) in the MTC sample is close to the baseline level estimated using an *E. coli* RNA sample sequenced in the same lane (0.1%). This result indicates that the MTC-purification procedure specifically captures the encapsulated RNAs, and that the transcriptome of a targeted subpopulation can be successfully isolated using MTCs without contamination from non-encapsulated transcripts.

Finally, we demonstrated that MTCs provide stable transcriptome storage inside host cells. Many factors could contribute to loss of MTC fidelity over time: the protein capsules could disassemble and reassemble, RNA within MTCs could decay, and expression of MTC proteins may be leaky after recording. To quantify the combined effect of all these potential sources, we examined the maintenance of MTC-encapsulated RNAs before and after shifting their host cells into a different environment that alters their transcriptomes. We first briefly induced MTC expression for one hour by adding IPTG (Isopropyl ß-D-1-thiogalactopyranoside) to Luria Broth. After washing off IPTG to shut off MTC expression, we introduced ethanol stress (4%) to the cells (Fig. 2d). The stress led to substantial remodeling of the host transcriptome, with 81 genes changed by >10-fold and an overall Pearson correlation coefficient of R=0.79 between mRNA levels pre- and post-stress (Fig. 2e). By contrast, the MTC samples collected before and after stress have almost identical RNA levels on a gene-by-gene basis, with an overall Pearson correlation coefficient of R=0.99 (Fig. 2e). The most extremely shifted gene in the host cell, *tnaA*, changed by 124-fold, whereas this same gene only changed by 1.5-fold across the MTC samples. These results demonstrate that MTC contents remain static despite large contextual changes in the cell state.

In summary, we established that MTCs can capture and preserve high-fidelity snapshots of the transcriptome in living cells. This approach opens several avenues of future applications. First, it will help elucidate the gene expression heterogeneities that precede distinct phenotypic outcomes during development, cellular differentiation, or stress survival. To do so, cells carrying pre-assembled MTCs can be sorted based on their final phenotypes, allowing their prior transcriptomes to be analyzed. This is facilitated by the fact that MTC-records are physically maintained inside the cell lineage^25^. MTCs can therefore nominate candidate genes for targeted studies. Second, MTCs can help elucidate cellular states adopted in hard-to-access locations, where in situ sample collection is difficult. The comparatively short recording time will enable precise capture of transitory responses, such as bacterial cells at specific locations inside an animal^21^. As MTC formation simply requires the self-assembly of two protein subunits, we anticipate that it will be generalizable to new systems, both eukaryotic and prokaryotic.

## Methods

### Strains

*B subtilis* strains were taken directly from *Bacillus subtilis* subsp. Subtilis str. 168. *E. coli* strains were generated from *Escherichia coli*, K-12 MG1655. This includes our wild-type control, which was taken directly from *Escherichia coli*, K-12 MG1655 and an MTC-containing strain which consists of *Escherichia coli*, K-12 MG1655 with the MTC-containing plasmid (described below). The plasmid was constructed using the method described below and transformed first into DH5α cells before being transformed into competent MG1655 cells using the method described below.

### Plasmids

pMP026 is a Kanamycin (Kan) resistant plasmid that contains constitutively produced LacI and IPTG-inducible MTC. It has pMB1 origin of replication. The full plasmid map can be found on our GitHub. This plasmid was constructed using the method described below. It was transformed into the *E. coli* strain K-12 MG1655 using the method outlined in greater detail below. This pMP026-containing *E. coli* strain was the one used for all experiments in this work.

### Plasmid Assembly

Plasmid components were either amplified using PCR from a pre-existing plasmid, followed by DpnI (NEB #R0176L) digestion, or were ordered directly as gene blocks from IDT. Plasmids were constructed by Gibson assembly, using the protocol and reagents associated with NEB #E2611 with a total volume of 4 µL. Plasmids were then transformed into Zymo Mix & Go DH5α Competent Cells (Zymo T3007) for verification and amplification before re-purification and transformation into MG1655 *E. coli*. All plasmid purification was done using overnight culture and the Zymo Zyppy Plasmid Miniprep Kit (Genesee 11-30). Plasmids were verified using Sanger sequencing (performed by Quintara), or by whole plasmid sequencing (performed by Plasmidsaurus). The plasmid associated with this work: pMP026, will be deposited to Addgene.

### Media

All experiments were conducted using Luria Broth (LB).

### Transformation

For transformation into Zymo Mix & Go DH5α competent cells (Zymo T3007), 100 µL of cells were thawed on ice per reaction. Then 1-4 µL plasmid DNA was added followed by gentle mixing. After 5-minute incubation on ice cells, 400 µL of prewarmed SOB medium was added, and the mixture was incubated in an Eppendorf tube for 1 hr at 37 °C with 300 rpm shaking. The mixture was then spread on pre-warmed agar plates with the appropriate antibiotic (50 µg/mL for MTC-plasmid-containing cells).

For transformation into non-competent wild-type MG1655 *E. coli*, the cells were made competent using the protocol described by Chung et al.^26^. 3 mL of LB was inoculated with a colony from a fresh agar plate. Cells were incubated at 37 °C 230 rpm for 1.5 to 2 hrs. 200 µL of cells were added to 200 µL ice cold TSS buffer (LB broth containing 10% (wt/vol) polyethylene glycol, 5% (vol/vol) dimethyl sulfoxide, and 50 mM Mg2+ at pH 6.5) and 1 µL plasmid. Cells were vortexed and incubated on ice for 20 - 30 minutes. Then cells were incubated for 45 - 60 min at 37 °C on the thermomixer with 900 rpm shaking. Finally, cells were plated on the appropriate antibiotic (50 µg/mL for MTC-plasmid-containing cells).

### Assessing MTC reproducibility

(Experiment shown in Figure 1c, Extended Figures 3 & 4) MG1655 containing pMP026 was streaked on a 50 µg/mL Kan marker plate and left to grow overnight at 37 °C. The next day, 3 single colonies were selected from the plates and incubated in separate test tubes with 5 mL LB and 50 µg/mL Kan for 2 hrs. These cultures were then back-diluted to allow for 12 doublings before reaching OD 0.3 in 500 mL of pre-warmed LB with 1 mM IPTG and 50 µg/mL Kan. Shortly before reaching OD 0.3, both the lysate and MTC samples were harvested. MTC samples were harvested by splitting the volume into four 50mL Falcon Tubes and spinning them down for 10 min at 4000 rpm at 4 °C in an Eppendorf 5810R Centrifuge. The supernatant was discarded, and the cell pellets were frozen for future protein purification. The lysate sample was collected by placing 500 µL of culture directly into a prewarmed RNA extraction solution (see below) and proceeding with RNA extraction.

### Assessing the Contamination from bulk lysate

(Experiment shown in Figure 2a-c) - Two samples were harvested for this experiment - MG1655 containing pMP026 and Wild Type *B. subtilis* cells (BS168).

The MG1655 containing pMP026 cells were streaked from a glycerol stock onto an LB + 50 µg/mL Kan plate and grown overnight. A colony from this plate was selected the next day and added to 10mL of LB with 50 µg/mL Kan and grown overnight at 37 °C on a rotator drum spinning ∼ 225 rpm. This overnight culture was then diluted into a volume of 500 mL to achieve 12 doublings before reaching OD 0.3 in pre-warmed LB with 50 µg/mL Kan. The culture was grown at 37 °C on a shaker plate - shaking at ∼ 225 rpm. At OD 0.3 1mM IPTG was added. At OD 2 the culture was divided between 50mL Falcon tubes and spun for 10 minutes at 4000 rpm in a pre-chilled (4 °C) Eppendorf 5810R Centrifuge. The Supernatant was then discarded, and the cell pellets were flash frozen in liquid nitrogen and stored at -80 °C.

The *B. subtilis* cells (BS168) were inoculated directly from a glycerol stock into 5 mL LB, then grown for 2 hours at 37 °C on a rotator spinning ∼ 225 rpm. This culture was then diluted to achieve 12 doublings before reaching OD 0.3 in pre-warmed LB. At OD 2 the culture was divided between 50 mL Falcon tubes and spun for 10 minutes at 4000 rpm in an Eppendorf 5810R Centrifuge. The Supernatant was then discarded, and the cell pellets were flash frozen in liquid nitrogen and stored at -80 °C.

### Assessing the stability of encapsulated RNAs

(Experiment shown in Figure 2d-e) - MG1655 containing pMP026 was streaked on a 50 µg/mL Kan marker plate and left to grow overnight at 37 °C. The next day single colonies were selected from the plates and incubated in a test tube with 5 mL LB and 50 µg/mL Kan for 2 hrs. This culture was then back diluted in a 500 mL volume with 50 µg/mL Kan to achieve 10 doublings before reaching OD 0.3. Once the cultures reached OD 0.3 MTC recording was initiated by the addition of 1 mM IPTG. After 1 hr, the capsule induction was shut off by filtering and washing the cells. The filter was washed with half the original cell volume of prewarmed LB. The cells were then resuspended in the original volume of LB without any antibiotics or IPTG. Pre-Stress Lysate and MTC samples were collected 15 minutes post-wash. The MTC sample was collected by filtering 2 volumes of 125 mL of culture. The cells were then resuspended in 2 50 mL Falcon tubes in pre-chilled (4 °C) LB. These tubes were then centrifuged for 10 min at 4000 rpm at 4 °C in an Eppendorf 5810R Centrifuge. The supernatant was discarded, and the cell pellets were frozen for future protein purification. The lysate sample was collected by placing 500 µL of culture directly into a prewarmed RNA extraction solution (see below) and proceeding with RNA extraction. To the remaining 250 mL of culture, we added a 4% ethanol stress, and allowed the cells to grow for 45 minutes before harvesting the post-stress MTC and lysate samples in a manner similar to the one described above.

### Protein Purification

Harvested Cell Pellets were rehydrated in 40 mL Lysis Buffer (150 mM Imidazole, 250 mM NaCl, 25 mM Tris-HCL, pH 8, with Protease Inhibitors) and the OD 600 values of each rehydrated solution were measured. Each sample was then sonicated using the 450W Ultrasonic Homogenizer (10 to 300 mL) from US Solid following a program of 2 seconds on at 25% power, followed by a 6 second off break for a total of 10 minutes. Lysate was then clarified by centrifugation at 4 °C using a Sorvall RC-5B Refrigerated Superspeed Centrifuge at 10,000 rpm for 45 min, after which the clarified supernatant was retained for loading on the protein column. To prepare the protein column we loaded 2 mL (1 mL column volume) of Nickel NTA resin onto the 5 mL Polypropylene columns from Qiagen and the resin buffer was allowed to drain. The resin was then rinsed with 15 column volumes of ddH2O followed by 15 column volumes of wash buffer (150 mM Imidazole, 250 mM NaCl, 25 mM Tris-HCL, pH 8) to equilibrate the column. After this the clarified supernatant is loaded onto the column. The column is then washed with 15 column volumes of the wash buffer followed by an elution with the elution buffer (500 mM Imidazole, 250 mM NaCl, 25 mM Tris-HCL, pH 8). 3 column volumes of the elution buffer were used, but only the last 2 column volumes were kept. The purified proteins are then treated with RNase A (1 µL/mL 20 °C for 10 minutes) before proceeding with RNA extraction.

### RNA Extraction

Add a prewarmed (65 °C) solution of 500 mL phenol acid chloroform with 29 µL 20% SDS to 500 mL of sample to a 1.5 mL Eppendorf in a thermomixer (split between multiple tubes if needed). Incubate at 65 °C for 5 minutes at 1,400 rpm, followed by a 5-minute incubation of the samples on ice. Spin the samples at 20,000 g for 2 minutes then transfer the top aqueous layer to a new non-stick tube. Add 45 µL 3M NaAc (pH 5.4), 500 mL 100% Isopropanol and 1 µL GlycoBlue coprecipitant. Chill the tubes at -80 °C for 30 min and then spin at 20,000 g for 60 min at 4 °C. Remove and discard the supernatant then add 250 mL pre-chilled 80% Ethanol and spin at 20,000 g for 5 min. Remove and discard the supernatant then resuspend the samples in 100 µL 10 mM Tris 7. Process the sample using the modified version of the Zymo-5 RNA Clean and Concentrator columns to remove all RNAs < 200 nt (to remove tRNAs). Elute the sample in 85 µL DEPC-water and proceed immediately with DNase treatment.

### DNase Treatment

Add 10 µL 10x Turbo DNase buffer and 5 µL Turbo DNase to 85 µL RNA. Incubate for 20-30 min at 37 °C in Eppendorf thermomixer without shaking. Add 330 µL 100% Ethanol, 11 µL 3M NaAc (pH 5.4) and 1 µL GlycoBlue coprecipitant. Chill the tubes at -80 °C for 30 min+ (possible stopping point) and then spin at 20,000G for 60 min at 4C. Remove and discard the supernatant then add 250 mL pre-chilled 80% Ethanol and spin at 20,000 g for 5 min. Remove and discard the supernatant then resuspend the samples in 13.5 µL DEPC-water. Measure concentrations using High Sensitivity RNA kit for Qubit, then proceed with library preparation.

### Library Preparation

Libraries in this paper were prepared using NEB’s rRNA Depletion Kit (Bacteria) for rRNA removal with beads (NEB #E7860). Post rRNA removal samples were prepared for either Illumina or Singular Sequencing using NEBNext Ultra II RNA Library Prep Kit for Illumina with Beads (NEB#E7775). Singular specific primers for the libraries sequencing on Singular which had the S1/S2 handles instead of Illumina’s p7/p5 handles.

### Code Availability

Raw sequencing fastq files were trimmed using seqtk and cutadapt to remove bases of low quality and adapters. Reads were then aligned using bowtie (version 1)^27^, after which the density of the 5’ ends was quantified using SAMtools^28^and the CDS files for each genome were used to quantify how many transcripts were found within each gene. The complete scripts for raw analysis, as well as processed data files (in the format of counts per gene) can be found on our GitHub (https://github.com/gwlilabmit/MTC_2023_Scripts). Jupyter notebooks also exist for each of the plotted subfigures and extended figures.

### Data Availability

Raw fastq files associated with this work have been deposited to the NCBI’s SRA database. This submission can be previewed using this reviewer link: (https://dataview.ncbi.nlm.nih.gov/object/PRJNA1024409?reviewer=bcc60b8vat29csnmgjv5fn12ba). They are associated with BioProject SUB13877560 and will be made public at the time of publication or submission to BioRxiv. Processed data files and figure source data can be found on our GitHub (https://github.com/gwlilabmit/MTC_2023_Scripts), Non-sequencing data (i.e. the doubling rate data in Extended Figure 2) can also be found in the GitHub.

## Acknowledgements

We would like to thank Lydia Herzel, Matthew Tien, and Robert Battaglia for all their help with various protocols used in this work. We would also like to thank Katherine (Julia) Dierksheide, James (Scott) McCain, and James Xue for their helpful discussions and suggestions for framing of this work. Julian Stanley, Jenny Cascino, Manraj Gill, Andrew Savinov, Mandy Levine, Hannah LeBlanc, James Taggart, Grace Johnson, and Cassandra Burgos-Schaening also provided useful feedback, suggestions, ideas, and criticism over the course of many group meetings and presentations. Furthermore, we would like to thank Joseph (Joey) Davis and Paul Blainey for their expert feedback and suggestions.

This work was made possible in collaboration with the MIT Biology Bio Micro Center, and in particular, we are grateful to Stuart Levine, Christopher Hallee, and Noelani Kamelamela for facilitating the sequencing and qPCR work reported in this paper.

This work was supported by NIH grant R35GM124732, the NSF CAREER Award MCB-1844668, the Smith Odyssey Award, and the Pew Biomedical Scholars Program.

## Author Contributions

M.P., G.L., and D.M. conceived of the presented idea and generated a candidate list of molecules to test as MTCs. M.P. and J.R. conducted the experiments in this work. M.P. conducted analysis of experimental results. G.L. advised on experimental design and data analysis, as well as suggested relevant literature. G.L. and M.P. wrote the final manuscript. All authors discussed results and commented on the manuscript.

