## Extended Figures for "Molecular Time Capsules Enable Transcriptomic Recording in Living Cells"

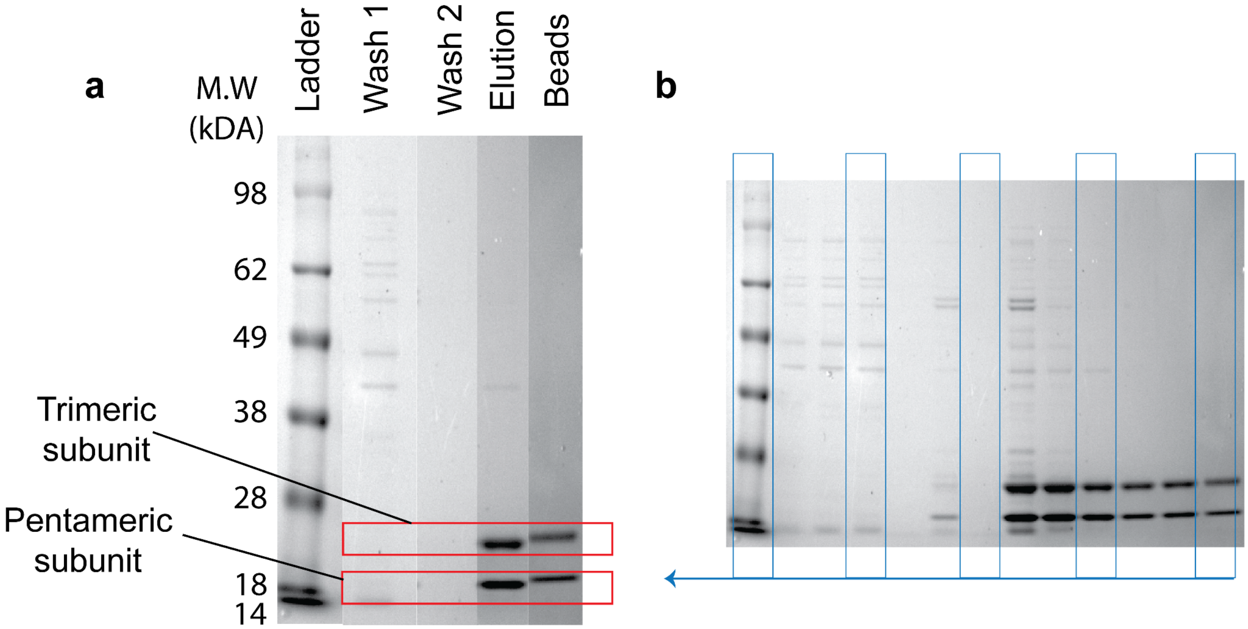


**Extended Figure 1: Protein Purification of MTCs.** MTC protein complexes can be eluted without substantial detected contamination. **a,** Samples of the flow-through from the affinity purification process used to purify the MTCs (see the methods section for more detail) run on an SDS PAGE gel. The ladder column corresponds to the See-Blue prestained protein ladder from Invitrogen. Wash 1 represents the flow through from the first wash of the column after loading the supernatant. Wash 2 represents the flow through of the second wash. The Elution column represents the eluted MTCs. The final column, Beads, corresponds to a sample of the Nickel-NTA beads post-elution. The pentameric and trimeric subunits are highlighted. **b,** The full MOPS SDS-Page gel that the lanes from subfigure **a** were extracted from. The columns extracted to make sub-figure **a** are highlighted in blue. Other columns correspond to other purification conditions which were not used in this work.

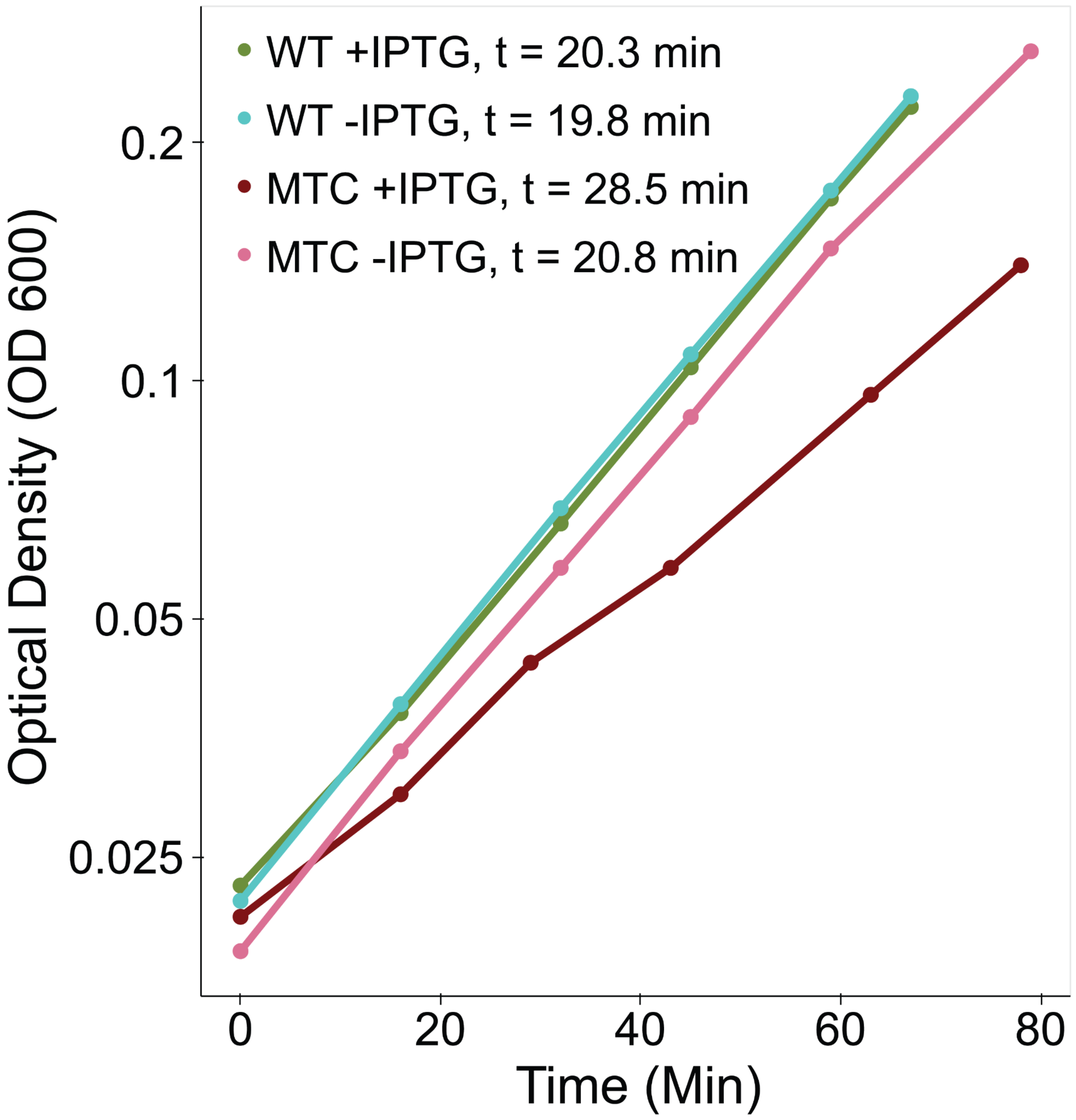


**Extended Figure 2: Impact of Extended Induction of MTCs on Doubling Time**. This figure contains a semi-log plot representing the optical density (measured at 600 nm) of bacterial cultures over time (minutes). WT corresponds to MG1655 *E. coli* (no vector), while MTC refers to MG1655 *E. coli* that carry pMP026 (the MTC plasmid). +IPTG samples had 1 mM IPTG added at the moment of back-dilution from overnight culture. -IPTG samples were back diluted into luria broth without the addition of any IPTG. MTC samples had 50 µg/mL of Kan added to each culture at the moment of back-dilution. The “t” in the legend above corresponds to the calculated doubling time in minutes for each sample. The +IPTG MTC sample has a doubling time of 28.5 minutes, which is ~ 8.5 minutes slower than WT samples.

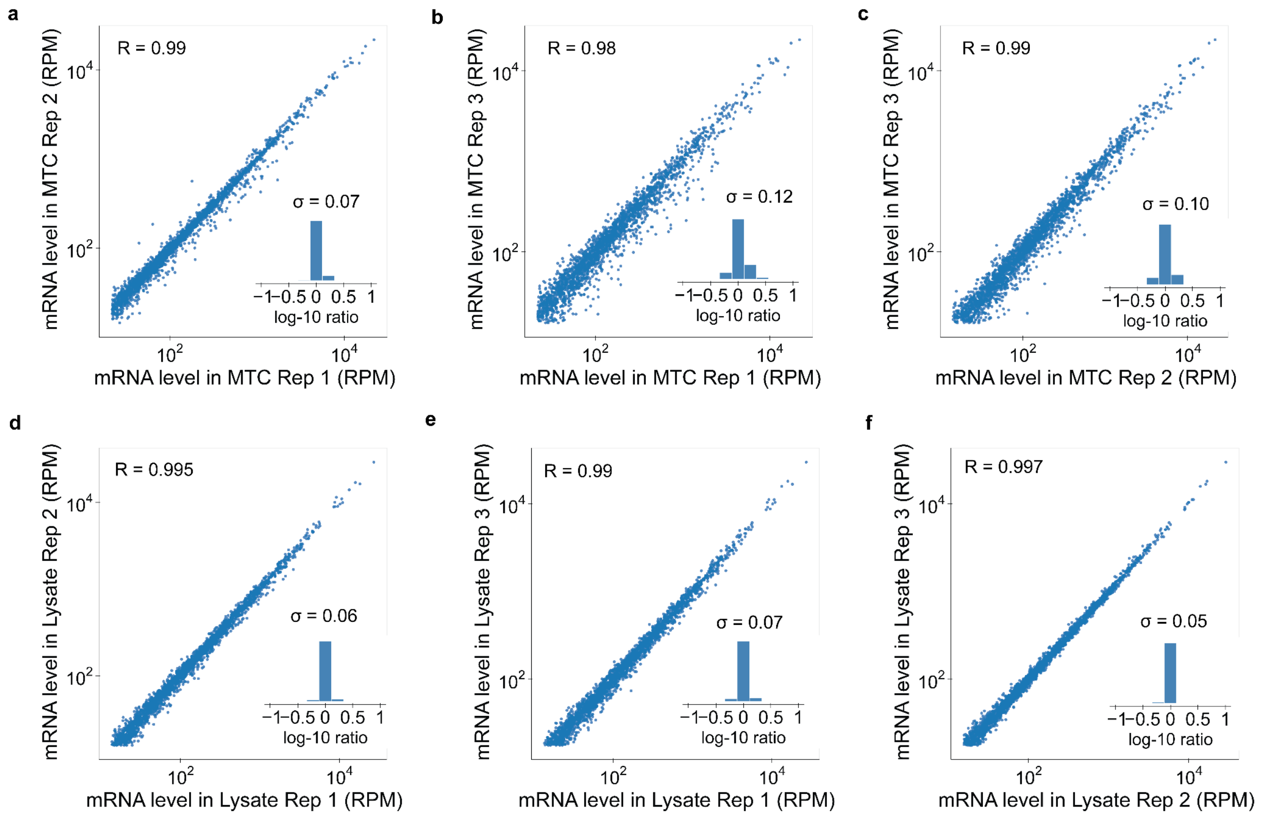


**Extended Figure 3**: **Reproducibility of MTC-capture and of final lysate across 3 biological replicates.** Subfigures **a-c** plot a gene-by-gene read per million (RPM) scatterplot comparing MTC-encapsulated mRNAs across 3 biological replicates (between replicates 1 and 2 R = 0.993 (n = 2302) the **σ** of fold-change distribution = 1.2 fold; 1 and 3 R = 0.976 (n = 2277) the **σ** of fold-change distribution = 1.3 fold; 2 and 3 R = 0.985 (n = 2395)) the **σ** of fold-change distribution = 1.3 fold. **d-f** demonstrates the corresponding gene-by-gene RPM values of mRNAs taken from the general lysate (between replicates 1 and 2 R = 0.995 (n = 2430) the **σ** of fold-change distribution = 1.2; 1 and 3 R = 0.994 (n = 2374) the **σ** of fold-change distribution = 1.2; 2 and 3 R = 0.997 (n = 2364) the **σ** of fold-change distribution = 1.1). All Pearson correlation coefficients (R) are calculated on log-transformed values (as is plotted) for reads with > 100 sequencing reads. The “n” reported for each Pearson correlation coefficient corresponds to the number of genes used in that calculation.

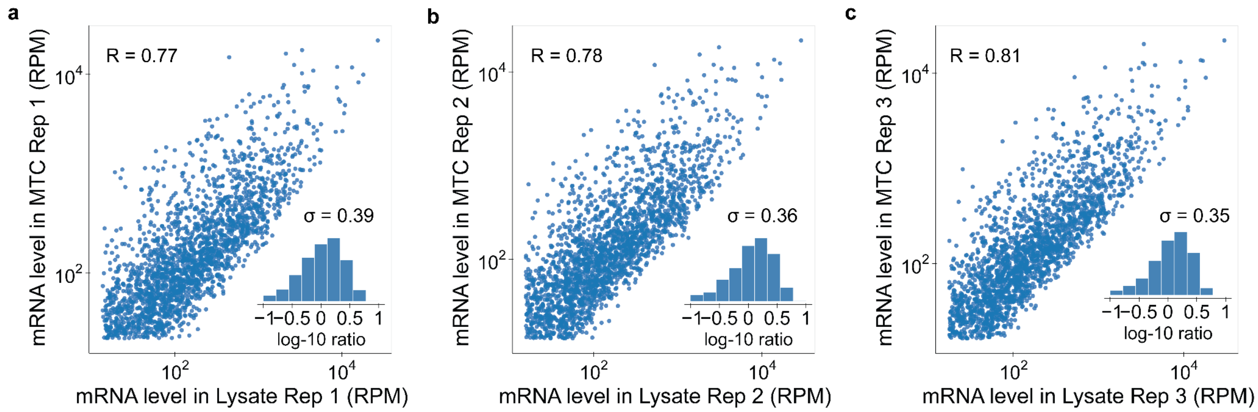


**Extended Figure 4: Similarity between MTC-encapsulated transcripts and cellular transcriptome across 3 biological replicates. a-c** Plot three biological replicates of MTC-encapsulated mRNAs and lysate purified mRNAs plotted on a gene-by-gene basis with read per million (RPM) value for each sample (between replicates 1 and 2 R = 0.766 (n = 2147) the **σ** of fold-change distribution = 2.5 fold; 1 and 3 R = 0.783 (n = 2235) the **σ** of fold-change distribution = 2.5 fold; 2 and 3 R = 0.813 (n = 2168)) the **σ** of fold-change distribution = 2.5 fold. While this represents a good agreement between MTC-encapsulated RNAs and lysate RNAs, it is important to note that the variance which contributes to the spread is highly reproducible - as is demonstrated in Extended Figure 3. This high reproducibility means that in using the MTCs it is best to compare MTCs between two conditions or populations of interest to detect potential up-regulated or downregulated genes rather than looking at the MTC versus lysate RPM values. All Pearson correlation coefficients (R) are calculated on log-transformed values (as is plotted) for reads with > 100 sequencing reads. The “n” reported for each Pearson correlation coefficient corresponds to the number of genes used in that calculation.

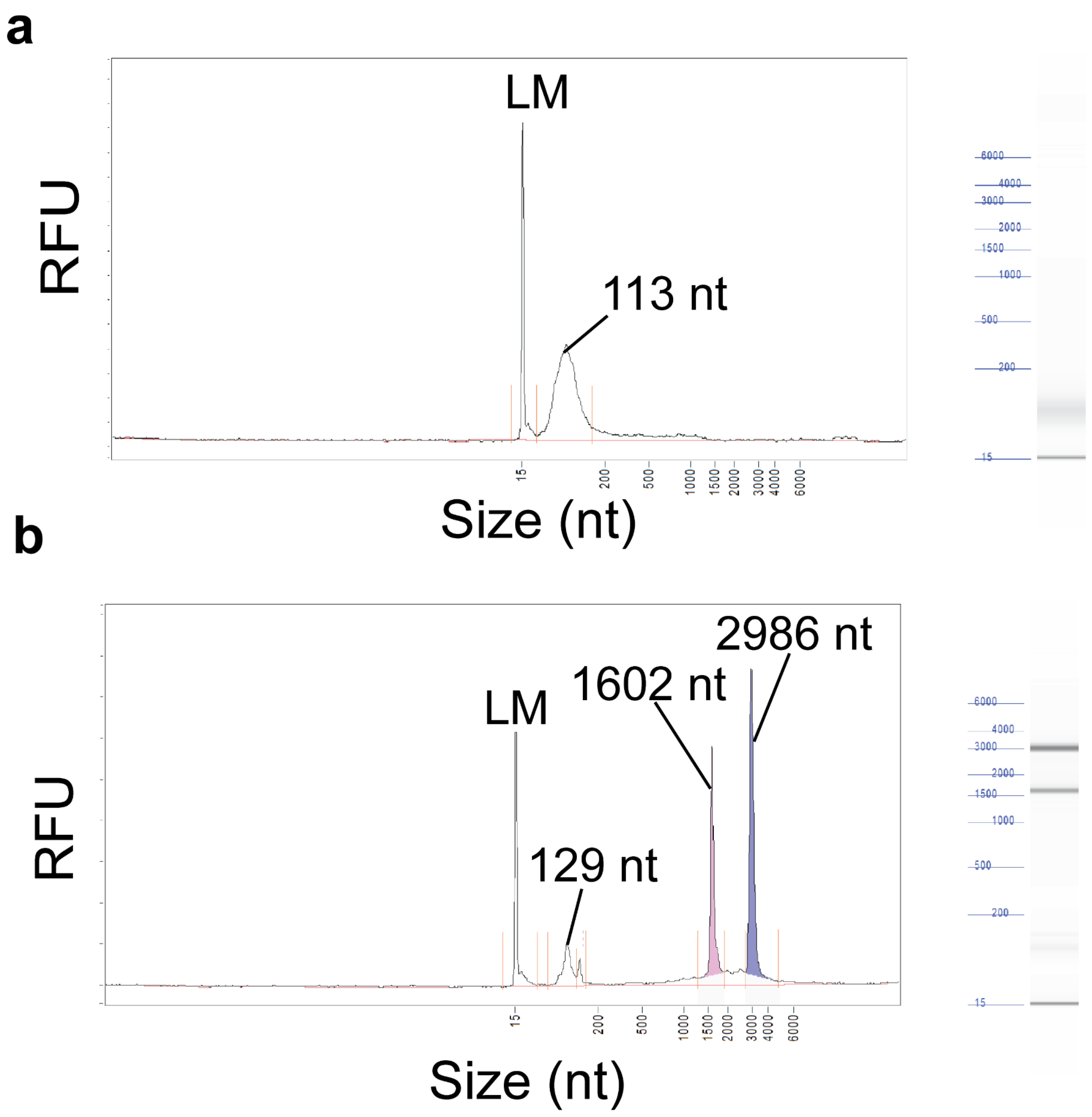


**Extended Figure 5 - Length Distribution of MTC Encapsulated RNAs in Comparison to General Lysate RNAs.** Shown here is the output of the length distribution measured via Fragment Analyzer. **a**, The length distribution of MTC-encapsulated transcripts. The distribution is dominated by very short transcripts that peaks at 113 nt. **b**, The length distribution of the general lysate. Highlighted are the peaks at ~130 nt, ~16k nt and ~30k nt which we believe correspond to rRNAs. In both plots LM refers to the lower molecular weight standard used by the Fragment Analyzer.
